# A zebrafish model for studying mechanisms of newborn hyperbilirubinemia and bilirubin induced neurological damage

**DOI:** 10.1101/2023.07.26.550752

**Authors:** Metehan Guzelkaya, Ebru Onal, Emine Gelinci, Abdullah Kumral, Gulcin Cakan-Akdogan

## Abstract

Unresolved neonatal hyperbilirubinemia may lead to accumulation of excess bilirubin in the body, and bilirubin in the neural tissues may induce toxicity. Bilirubin induced neurological damage (BIND) can result in acute or chronic bilirubin encephalopathy, causing temporary or lasting neurological dysfunction or severe damage resulting in infant death. Although serum bilirubin levels are used as an indication of severity, known and unknown individual differences affect the severity of the symptoms. The mechanisms of BIND have not been fully understood yet. Here, a zebrafish newborn hyperbilirubinemia model is developed and characterized. Direct exposure to excess bilirubin induced dose and time dependent toxicity linked to the accumulation of bilirubin in the body and brain. Introduced bilirubin was processed by liver which increased the tolerance of larvae. BIND in larvae was demonstrated by morphometric measurements, histopathological analyses and functional tests. The larvae that survived hyperbilirubinemia displayed mild or severe morphologies associated with defects in eye movements, body posture and swimming problems. Interestingly, the plethora of mild to severe clinical symptoms were reproduced in the zebrafish model.

**Summary statement:** This alternative newborn hyperbilirubinemia model in zebrafish, reports detailed analyses of bilirubin toxicity, recovery, and bilirubin induced neurological damage in varying degrees. Various clinical symptoms of BIND is successfully reproduced.

## 1. INTRODUCTION

Bilirubin is the end product of hemoglobin breakdown, a beneficial molecule at low levels due to its antioxidant properties, but becomes toxic when elevated due to its insoluble nature (1). Unconjugated bilirubin (UCB) is carried by serum albumin protein to the liver where it is conjugated to glucuronic acid by the hepatic UDP glucuronosyltransferase 1A1 (UGT1A1), and produced bilirubin-glucronidic acid conjugates are secreted into bile. Increased erythrocyte turnover which leads to increased bilirubin production, and the delayed activation of UGT1A1 in newborns can lead to hyperbilirubinemia (2, 3). Jaundice is the mild form of neonatal hyperbilirubinemia which occurs in 60% of term neonates and almost all pre-term newborns, and often self-resolves within days (3). However, infants may develop severe hyperbilirubinemia when serum bilirubin levels increase above 20 mg/dL, and UCB can accumulate in tissues including the brain (4). UCB accumulation in the central nervous system can lead to bilirubin-induced neurological damage (BIND) (5). To prevent BIND, infants with significant hyperbilirubinemia are first treated with phototherapy which increases the amount of soluble bilirubin photo isomers. Next, blood transfusion is performed to eliminate excess bilirubin from the serum (6, 7). BIND may result in acute bilirubin encephalopathy (ABE) or chronic bilirubin encephalopathy (CBE, aka kernicterus) that leaves permanent neurological damage characterized by oculomotor problems, auditory neuropathy, muscle tonus and movement anomalies (8). Individual factors such as serum albumin levels and its bilirubin binding capacity, genetic susceptibilities, metabolic differences, factors that control central nervous system exposure to bilirubin and bilirubin clearance from CNS are thought to affect the progression of the disorder (9).

Although some animal models are developed, the molecular mechanisms of BIND in infants and the predisposing factors are not fully understood. The *ugt1* mutant rat (Gunn Rat) is a genetic model with lifelong hyperbilirubinemia and a good model to simulate Criegler-Najjar syndrome, however these animals do not develop hyperbilirubinemia at the newborn stage (10). Another genetic model is generated in mice, which induces neonatal lethality (11). Induction of hemolysis by drug administration or direct injection of bilirubin into the brain are among methods to induce newborn hyperbilirubinemia (12-14). Each model has its own advantages and disadvantages, as well as technical challenges; and need for in vivo models that will allow mechanistic studies to be performed for understanding BIND mechanisms and develop preventive strategies is not fulfilled (8, 15).

Zebrafish model can offer advantages such as ease of live imaging throughout embryonic and larval stages, ease of chemical exposure and genetic modifications. However, no zebrafish hyperbilirubinemia model was developed to date, although the bilirubin metabolism seems to be conserved in zebrafish. Zebrafish produce hemoglobin as of 48 <u>h</u>ours <u>p</u>ost fertilization (hpf) and *heme oxygenase* and *biliverdin reductase* genes are expressed at this stage (16). The transition from embryo to free-feeding larvae is completed by 120 hpf, while gastrointestinal system and liver gain full functionality, and liver UGT enzymes are expressed (17-20). Based on this knowledge, we hypothesized that the newly hatched larvae at 48 hpf resembles a newborn baby in terms of bilirubin metabolism and liver function and the window for neonatal hyperbilirubinemia induction should be between 2 - 5 dpf. Accordingly, hyperbilirubinemia was induced by direct exposure of early larvae to bilirubin. The conditions of acute hyperbilirubinemia induction, tissues of bilirubin accumulation, routes of bilirubin elimination, histopathology of diseased larvae were examined. Finally, lasting neurological effects of acute hyperbilirubinemia were examined in recovered larvae. Several clinical symptoms were successfully reproduced in the zebrafish neonatal hyperbilirubinemia model.

## 2. MATERIALS AND METHODS

### Zebrafish maintenance

Zebrafish were reared under standard conditions, at 28°C, under 14/10h light/dark cycle, at Izmir Biomedicine and Genome (IBG) Center Zebrafish Vivarium. The Casper zebrafish line (roy^-/-^; nacre^-/-^) embryos were incubated at 28°C in E3 (5 mM NaCl, 0.17 mM KCl, 0.33 mM CaCl_2_·2H_2_O and 0.33 mM MgCl_2_·6H_2_O, 1% methylene blue) medium, before and during the experiment (21). All animal procedure was in compliance with Directive 2010/63/EU and was approved by IBG Local Ethics Committee for Animal Experimentation (IBG-HADYEK), with protocol number 2021-012.

### Bilirubin treatment

85.5 mM bilirubin stock was prepared in 0.25 M NaOH, 6 mM Bovine Serum Albumin (BSA) stock was prepared in PBS. Bilirubin: BSA (3:1) solution containing 900 μM bilirubin was prepared by diluting BSA in PBS and adding was by adding required volume of 85.5 mM bilirubin. This intermediate stock was aliquoted and frozen, fresh dilutions were made in E3 at the time of treatment. Dechorionated 48 hpf larvae were treated in 24-well plates, protected from the light and incubated at 28°C. At the end of exposure, larvae were washed with E3 three times to remove excess bilirubin and to take images.

### Imaging and quantitation

Larvae were imaged live (anesthetized with tricaine) or after fixation with 4% formaldehyde. Samples were embedded in 1% low melting agarose and mounted on glass bottom petri dishes. Olympus SZX10 stereomicroscope and Zeiss LSM880 confocal microscope were used for imaging. Sizes of larval structures and relative amount of accumulated bilirubin was measured with ImageJ software using brightfield stereomicroscope recorded dorsal images of the larvae. For bilirubin quantitation the RGB images were converted to gray scale with ImageJ (version 1.53q), and inverted images were used to measure the mean gray value (MGV) in defined areas.

### Spectrophotometric Analysis

60 larvae were crushed in 80 μL of PBS, debris was removed with centrifugation at 6000 rpm, for 4 minutes. 20 μL of 0.25 M NaOH solution was gently pipetted onto the supernatants to solubilize bilirubin. The mixture was centrifuged for 2 minutes at 6000 rpm. 70 μL of supernatants were transferred to black 96-well plates, and the absorbance spectrum was recorded at 400 - 500 nm using a spectrophotometer, peak of absorbance was found to be at 420 nm. Bilirubin standard curve was obtained by measuring spectra of 0.01, 0.03, 0.05, 0.1, 0.2, 0.3, 0.4, 0.5 and 0.8 mg/dL bilirubin:BSA in control larval lysates. Formula (y = −6.1064x^2^ + 8.1667x – 0.0095; R^2^:0.99) obtained from the concentration/absorbance plot was used to calculate bilirubin concentration in lysates. Amount of average bilirubin retained in the larval body was reported as ng.

### Gaze limitation test

6 dpf zebrafish were placed in 3% methylcellulose without anesthesia, 30-second-long videos were recorded dorsally under stereomicroscope, while light flashes were applied 5 times with 6-second intervals. Turn angles were calculated with ImageJ.

### Quantitation of free swimming

Larvae were imaged in 24-well plates at 4 dpf and 6 dpf, with a stereomicroscope. 25 frame per second and 3-minute-long videos were recorded with SC50 camera. Trace lengths and speeds were analyzed using the Tracker 6.1.3 application auto tracker tool from Open-Source Physics (OSP).

### Statistical Analyses

Statistical significance analysis was done with GraphPad Prism software. Student’s T-Test was used for comparison of two groups, and One-way ANOVA was used for comparison of several groups to one control.

### Histopathological analysis

6 dpf zebrafish larvae were fixed with 10% neutral buffered formalin, overnight at 4^0^C and processed through a series of alcohol and xylene to embed in paraffin (22). 1 μm sections were obtained with a microtome. Hematoxylin and Eosin-stained sections were imaged (Olympus BX61).

## 3. RESULTS

### 3.1 Phenotypes of bilirubin exposed larvae

In order to induce hyperbilirubinemia in zebrafish, bilirubin ranging from 3 μM to 45 μM was added to the aqueous medium and the zebrafish larvae were treated for 24 hours or 48 hours, starting from 48 hpf. Effects of excess bilirubin on the developing zebrafish were monitored via imaging of the larvae every 24 h until 5 dpf (Fig.1 and Fig. S1). The yellow color of bilirubin which penetrated the zebrafish body was easily detected with brightfield stereomicroscope imaging.

Upon 24 h exposure between 2 - 3 dpf, 3 μM or 6 μM bilirubin caused yellow coloration of the head and accumulation in the otic vesicle, but did not induce any morphological defects or toxicity. After washout, gradual elimination of bilirubin from the body was observed at 4 dpf and 5 dpf. While the brain became less yellow, the gut accumulated more bilirubin. On the other hand, exposure to 9 μM and 15 μM bilirubin caused more prominent coloration of the whole body, with significant bilirubin deposition in the brain, otic vesicles and other tissues. Pericardial edema was observed at the end of 24 h exposure (Fig. 1D, F).

**Fig. 1.**
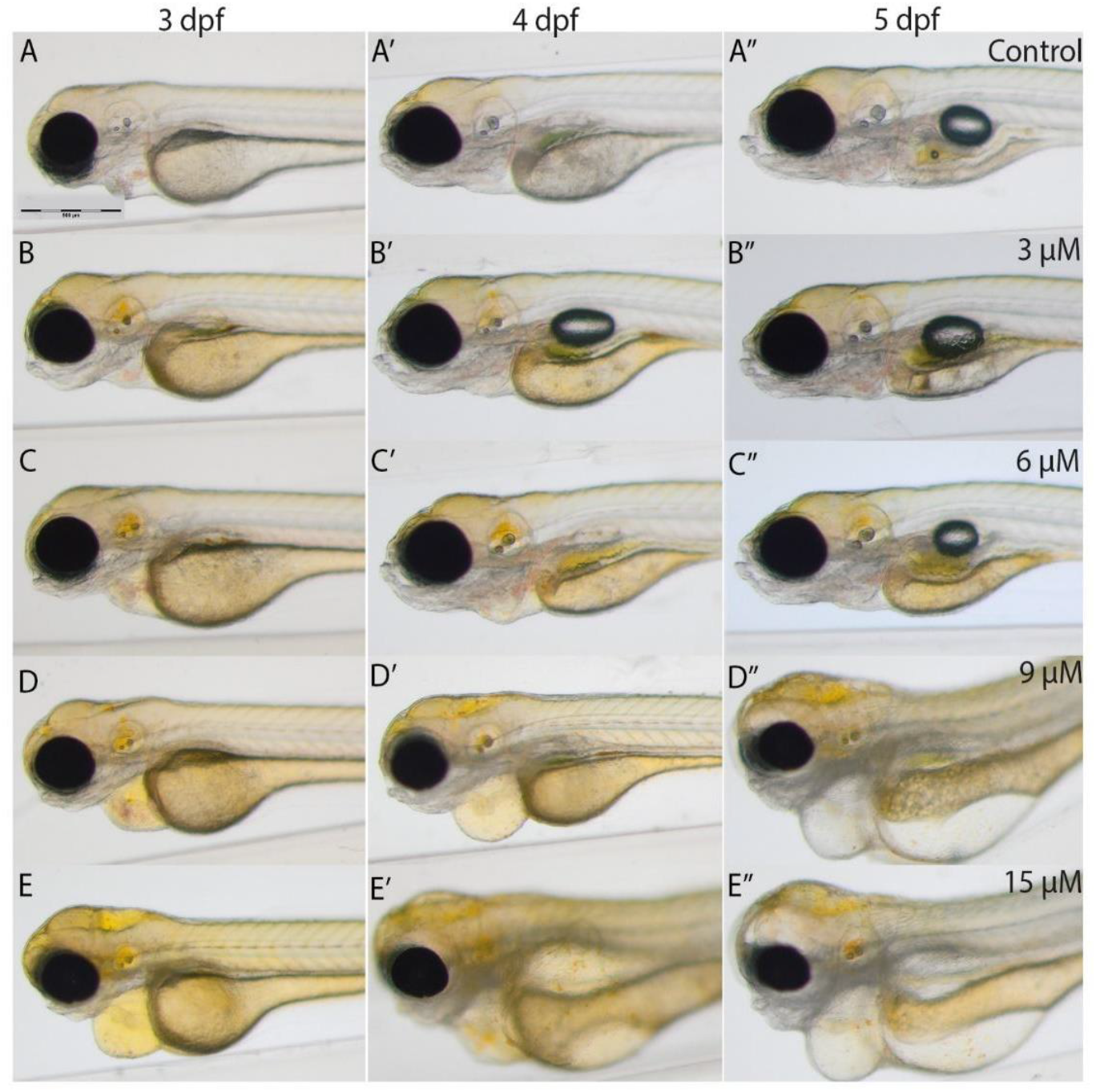
Images of bilirubin exposed larvae: Larvae were exposed to; **A-A’’**) carrier, **B-B’’**) 3 μM, **C-C’’**) 6 μM, **D-D’’**) 9 μM, and **E-E’’**) 15 μM bilirubin between 2 - 3 dpf. Representative images recorded at the end of treatment (3 dpf, left column), 1 day after washout (4 dpf, middle column) and 2 days after washout at 5 dpf (right column) are displayed in the figure. Scale bar: 200 μm, n=40

Treatment with 30 μM bilirubin caused lethality with full penetrance. 1 day after the washout, at 3 dpf, some bilirubin was detected in the gut area however the body was still yellow (Fig. 1D’, F’). The toxicity of 9 - 15 μM bilirubin was permanent; the treated larvae had heart and yolk edema, darkening of the brain and muscle structures at 4 dpf (Fig. 1D’, F’), and major deterioration of the larval body with smaller head, damaged gut, yolk edema and curved body was observed at 5 dpf (Fig. 1D’’, F’’).

Individual differences were observed among the treated larvae. While some larvae had a mild phenotype with heart edema, others displayed a severe phenotype with swelling around the eyes and in the head, major pericardial and yolk sac edema, distorted body posture, deteriorated muscles and loss of transparency (Fig. 2A, B). Toxicity induced by 9 - 15 μM bilirubin exposure for 24 h led to a significant shortening of the larval body as indicated by total length quantification of all treated larvae at 4 dpf (Fig. 2C). Tested lower doses did not affect the body length upon 24 h treatment while 48 h treatment with 6 μM bilirubin caused a mild but significant shortening of the body (Fig. 2C). Penetrance of severe phenotype increased with higher bilirubin dose. 34% of larvae which were treated with 9 μM between 2 - 3 dpf displayed a severe phenotype, whereas 15 μM bilirubin induced severe phenotype in nearly 90% of the larvae, and (Fig. 2D). From 4 to 7 dpf, progression of existing symptoms and an increase in the prevalence of severe phenotype was observed in all groups. Increase of exposure time to 48 h did not cause any toxicity at 3 or 6 μM doses, although a minor build-up of bilirubin in the body was observed (Fig. 2E, Fig S2). However, 9 μM bilirubin treatment for 48 h induced severe phenotype in 60% of the treated larvae and 15 μM led to severe phenotype which caused lethality in 90% of larvae by 5 dpf (Fig. 2E). Overall, these data showed that accumulation of bilirubin in zebrafish tissues and the damage induced by hyperbilirubinemia increased in a dose and exposure time dependent manner. Since clearance of bilirubin relies on liver, kidney and gut function, we hypothesized that if bilirubin exposure starts later during development, the bilirubin clearance would be more efficient resulting in less toxicity (23). As expected, when bilirubin exposure was applied between 3 - 4 dpf, the larvae tolerated higher doses of bilirubin, and incidence of severe phenotype was reduced (Fig. 2F).

**Fig. 2.**
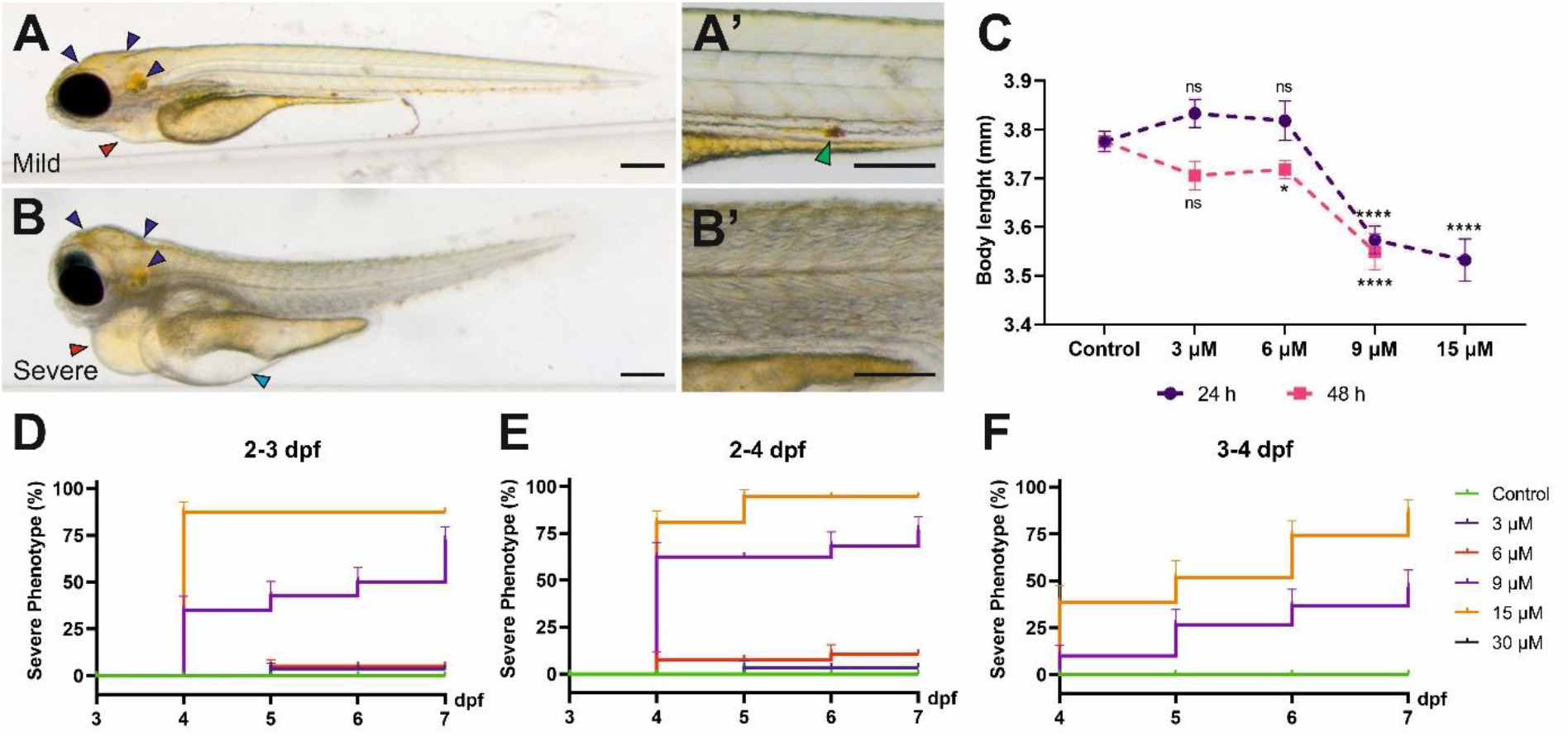
Mild and severe phenotypes: The larvae exposed to 9 μM bilirubin between 2 - 3 dpf, display **A)** mild or **B)** severe phenotype at 4 dpf. Close-up images are shown next to the overview images. Black, blue and red arrows represent bilirubin, yolk edema and pericardial edema, respectively. Green arrow indicates bilirubin excreted from the GI tract. Scale bars: 200 μm. **C)** Average body length of larvae treated with bilirubin, at 2 - 3 dpf for 24 h (purple line), or at 2 - 4 dpf for 48 h (orange line) shows a significant shortening of body length upon 9 - 15 μM bilirubin exposure. **D-F)** Penetrance of severe phenotype after exposure to bilirubin between **D)** 2 - 3 dpf, **E)** 2 - 4 dpf and **F)** 3 - 4 dpf. Larvae were treated with vehicle, 3 μM, 6 μM, 9 μM or 15 μM bilirubin. Mean ± SEM was calculated from triplicate treatment of 15 larvae in each group.

### 3.2 Bilirubin is processed by the liver and gut in zebrafish larvae

Histopathological analysis was performed to visualize bilirubin in tissues and investigate the effects of bilirubin on liver and gut which are organs involved in bilirubin metabolism and excretion. Since the development of the gastrointestinal system and liver is completed at the end of 5 dpf sections were taken on 6 dpf larvae that were exposed to 9 or 15 μM bilirubin between 2 −3 dpf or 3 - 4 dpf. Liver damage was visible in larvae that received bilirubin between 2 - 3 dpf (Fig. 3A-C). Liver vacuolation, disruption of hepatocyte tubules and enlargement of hepatic vessels were detected in the 9 μM group (Fig. 3B). Liver degeneration was more severe in the 15 μM group where large clusters of bilirubin were detected in disturbed bile ducts, large vacuoles, breakdown of hepatocytes, hepatocytes with enlarged or small nuclei were observed (Fig. 3C). The larvae that received 9 μM bilirubin treatment between 3 - 4 dpf had healthy livers with normal hepatocytes, hepatocyte tubules and general morphology (Fig. 3D). Detection of bilirubin in the liver also supports the hypothesis that bilirubin is metabolized in zebrafish larval liver and excreted into the bile. Although 15 μM bilirubin exposure between 3 - 4 dpf resulted in some toxicity, the livers of these larvae were healthier (Fig. 3E). The hepatocytic vacuoles were not observed, however the cytoplasm of the hepatocytes were basophilic, as in 9 μM 2 - 3 dpf treated group (Fig. 3B, E).

**Fig. 3.**
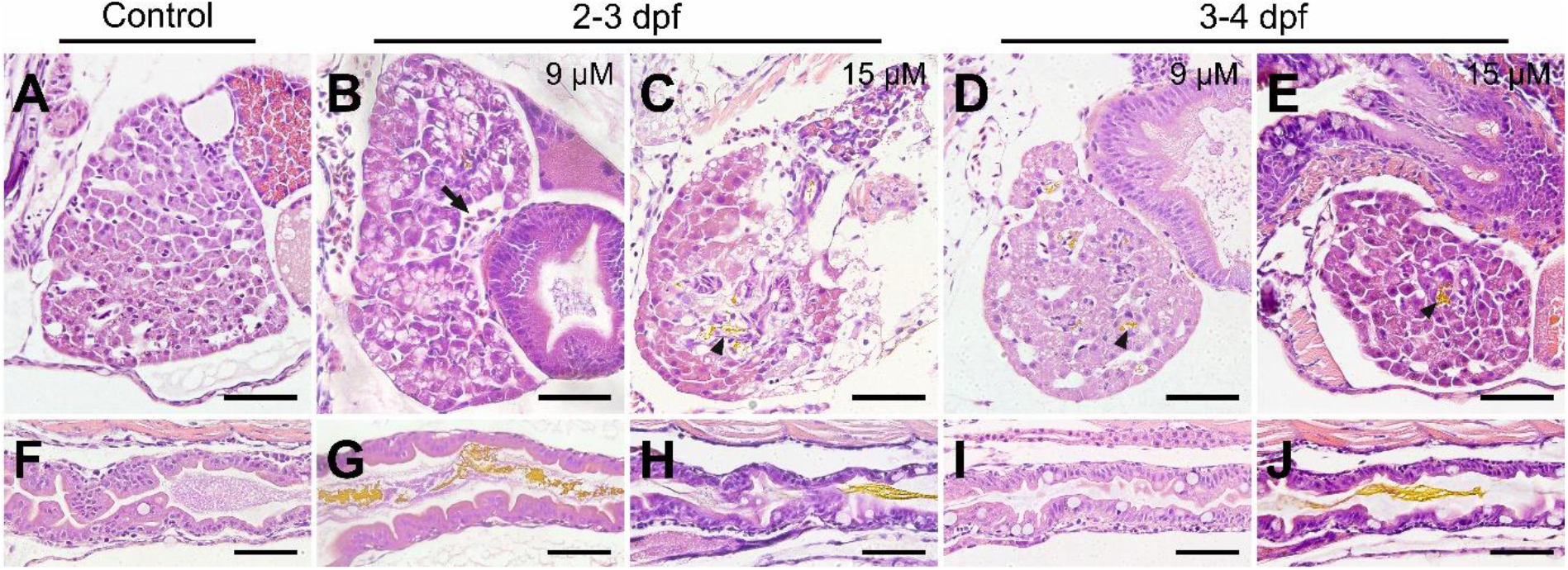
Histopathology of liver and gut after bilirubin exposure. **A-E)** Liver and **F-J)** gut tissues of 6 dpf larvae are stained with H & E after paraffin sectioning. **A, F)** Control larvae treated with vehicle. Larvae exposed to **B, G)** 9 μM, **C, H)** 15 μM bilirubin between 2 - 3 dpf. Larvae exposed to **D, I)** 9 μM, **E, J)** 15 μM bilirubin between 3 - 4 dpf. Arrow: enlarged vein, arrowhead: bilirubin in bile duct, scale bars: 50 μm.

Damage to gut was evident in 2 - 3 dpf exposed larvae, as disorganization of enterocytes, loss of gut folds and disruption of the mucosal layer when compared to control (Fig. 3F, G, H). Moreover, an increase of goblet cells was apparent. Later exposure to bilirubin was tolerated better in gut, especially at 9 μM dose the gut structure was not significantly affected, no bilirubin was detected in the lumen, although increase of goblet cell number was observed in some larvae (Fig. 3I). Finally, exposure to 15 μM bilirubin between 3 - 4 dpf affected the gut structure to the extent that folds were disturbed and some enterocytes were disorganized (Fig 3J).

### 3.3 Bilirubin accumulation in the larval body and brain

Total bilirubin retained in the body of the treated larvae was quantified spectrophotometrically. To this end, larval lysates were obtained, bilirubin was solubilized by addition of NaOH to the suspension and bilirubin absorbance was measured. The amount of retained bilirubin in the larval body was calculated in each experimental condition. Larvae exposed to 6 μM bilirubin for 24 h (2 - 3 dpf) accumulated around 780 ng of bilirubin in their body (Fig. 4A). 9 μM and 15 μM bilirubin exposure led to 1373 ng and 1676 ng bilirubin retention in the larval body, respectively. 48- hour exposure to low dose of 6 μM induced 1643 ng bilirubin accumulation, whereas maximum measured bilirubin accumulation, 2099 ng, was found in larvae treated with 9μM bilirubin for 48-hours.

**Fig. 4.**
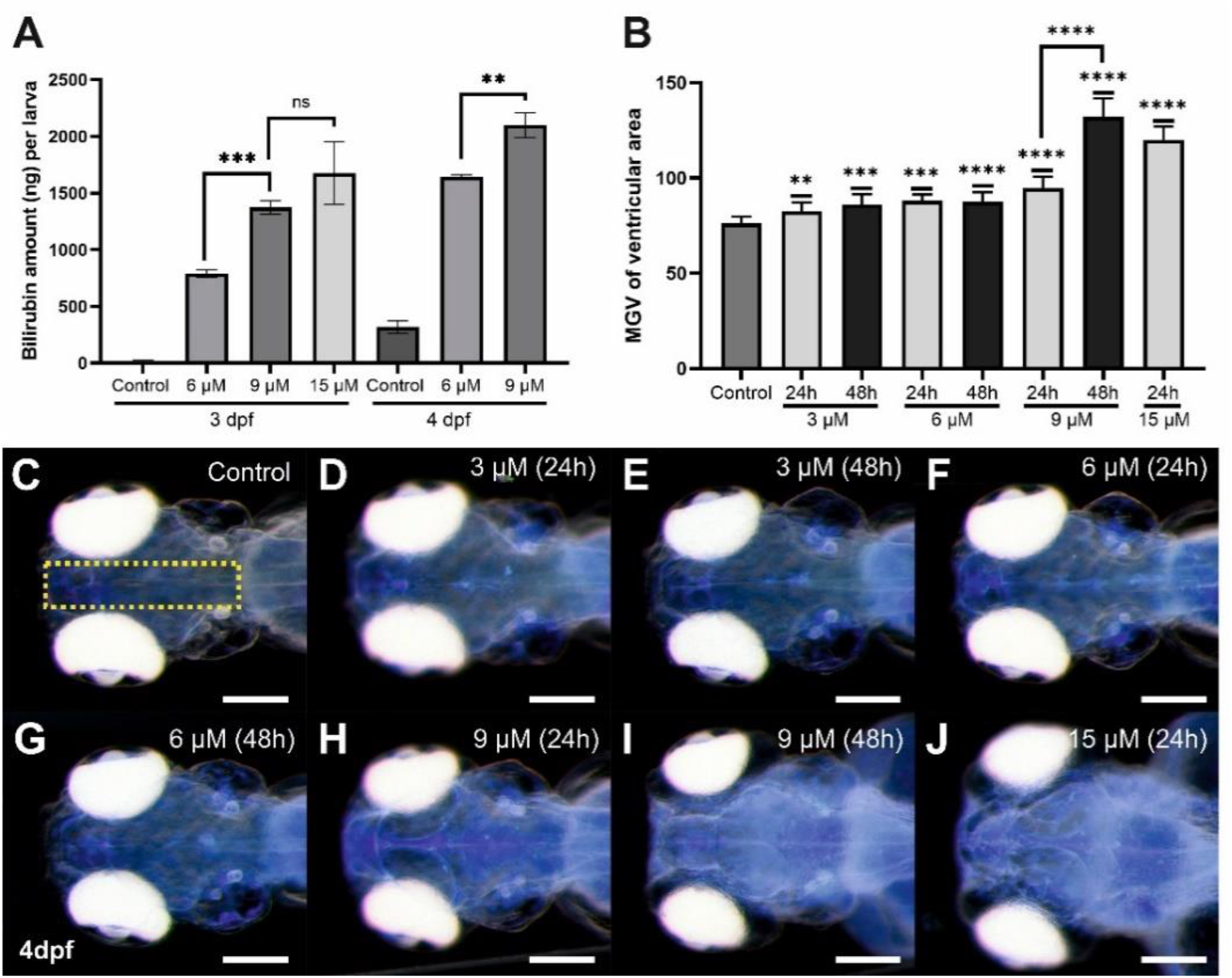
Quantitative analyses of bilirubin accumulation: **A)** Bilirubin concentration in larval body was measured from lysates, after exposure to 6 and 9 μM bilirubin for 24 or 48 h, mean values were plotted (n = 60 for each group, experiment was done in triplicate). **B)** Relative bilirubin accumulation in the brain at 4 dpf was plotted for each group (n = 15). **C)** Inverted images of the head which show bilirubin accumulates as bright blue spots were used for the graph in B. Scale bar: 200 μm

The spatial distribution of bilirubin depositions in the brain was observed easily when larval heads were imaged dorsally with a stereomicroscope at 4 dpf (Fig. S3). The depositions were most concentrated in the ventricles, and found to be spreading to a wider area as the dose and exposure time increased (Fig. S3). 9 or 15 μM bilirubin led to increased bilirubin staining throughout the brain, especially making the telencephalon and rhombencephalon readily distinguished (Fig. S3F-H). Accumulation of bilirubin in the brain was quantified by measuring the bilirubin signal at the middle of the brain images, and average signal intensities were plotted (Fig. 4B, C). A dose dependency was observed and maximum accumulation was detected after 24 h exposure to 15 μM. 48 h exposure to 3 or 6 μM bilirubin did not further increase the accumulation. On the other hand, 9 μM bilirubin exposure resulted in increased accumulation at 48 h (Fig. 4 B). Quantitation of bilirubin intensity in the brain was also performed at 3 dpf and 5 dpf and a similar trend was detected (Fig. S4). Similarly, bilirubin accumulation in the otic vesicle increased in a dose and time dependent manner (Fig. S5)

### 3.4 Hyperbilirubinemia caused neural damage in zebrafish larvae

Hyperbilirubinemia induced development of edema in the brain and around the eyes by 4 dpf, coupled to shrinking of these tissues (Fig. S3G-H). Eye size decreased and sclera of the ocular tissue was enlarged in a dose dependent manner (Fig. 5). Significant reduction of vertical and axial eye lengths was seen in both 9 μM and 15 μM bilirubin treated groups. Sclera enlargement was more prominent in 15 μM treated group with 20% increase in size. This enlargement was also observed at the back of the eyeball which caused an increase of the inter-ocular distance. On the other hand, brain tissue appeared smaller, and a significant dose dependent reduction of optic tectum was observed (Fig. 5E, Fig. S3).

**Fig. 5.**
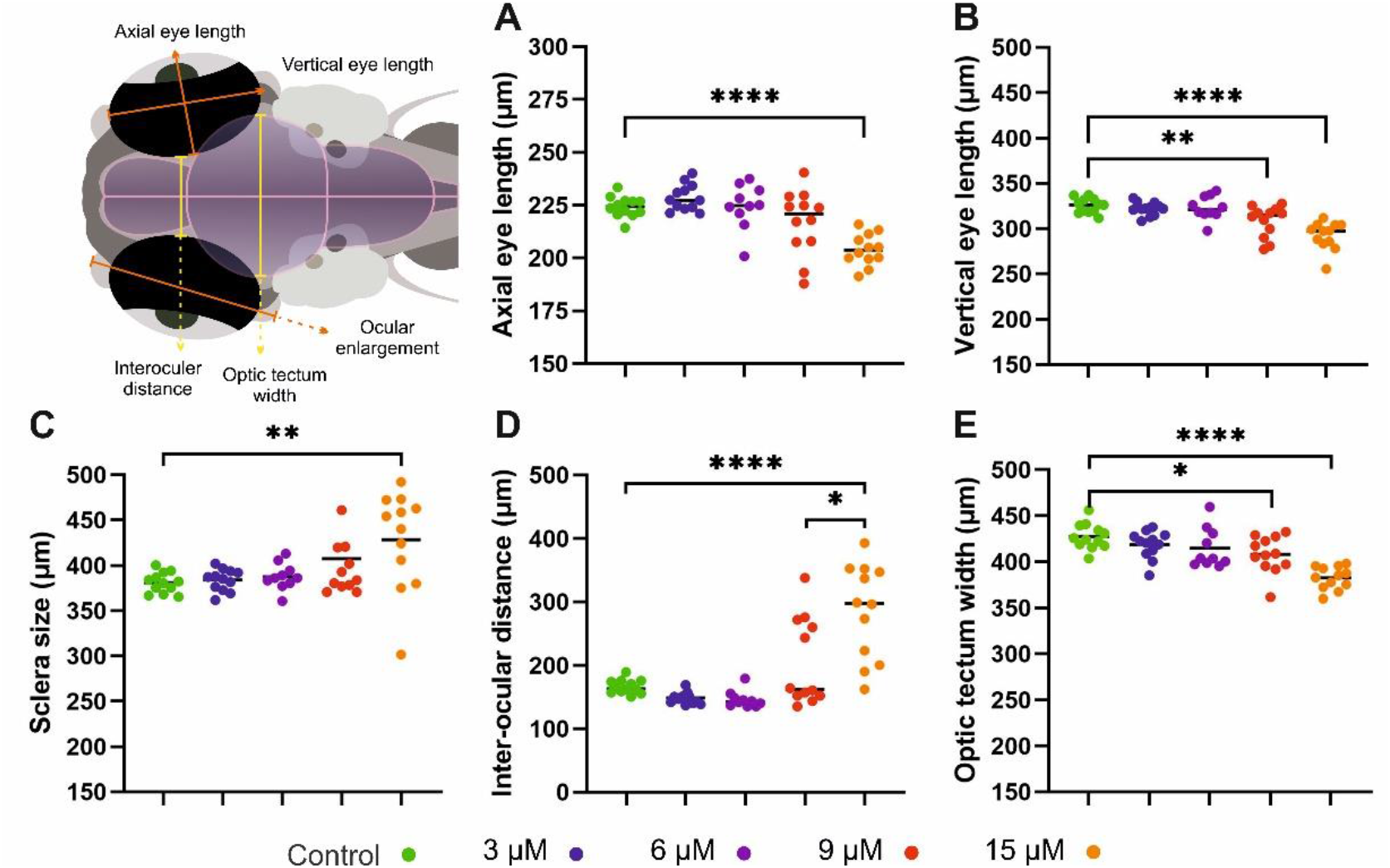
Morphologic analysis of neural tissues in bilirubin exposed larvae: Larvae exposed 3 μM, 6 μM, 9 μM and 15 μM bilirubin between 2 - 3 dpf were imaged at 4 dpf. **A)** axial eye length and **B)** vertical eye length decreased; **C)** sclera size and **D)** inter-ocular distance increased; **E)** optic tectum width decreased upon hyperbilirubinemia induction. (n=24, n=12, n=10, n=19 and n=13 for control group, 3 μM, 6 μM, 9 μM and 15 μM)

An increase in apoptosis was observed in larvae treated with 15 μM bilirubin (Fig. 6A). The degeneration in the brain tissue was visible on tissue sections of larvae, several days after washout, at 6 dpf (Fig. 6B - F). Bilirubin accumulation in the brain led to formation of edema and shrinking of the neuronal tissue in 9 μM exposed larvae (Fig. 6 C). Major acellular areas and fragmentation of the tissue was detected in the forebrain and hindbrain of 15 μM exposed larvae (Fig. 6D). These defects were less pronounced in the group that received bilirubin between 3 - 4 dpf (Fig. 6E-F). Otic vesicles were deformed in all groups regardless of the exposure time (Fig. S6).

**Fig. 6.**
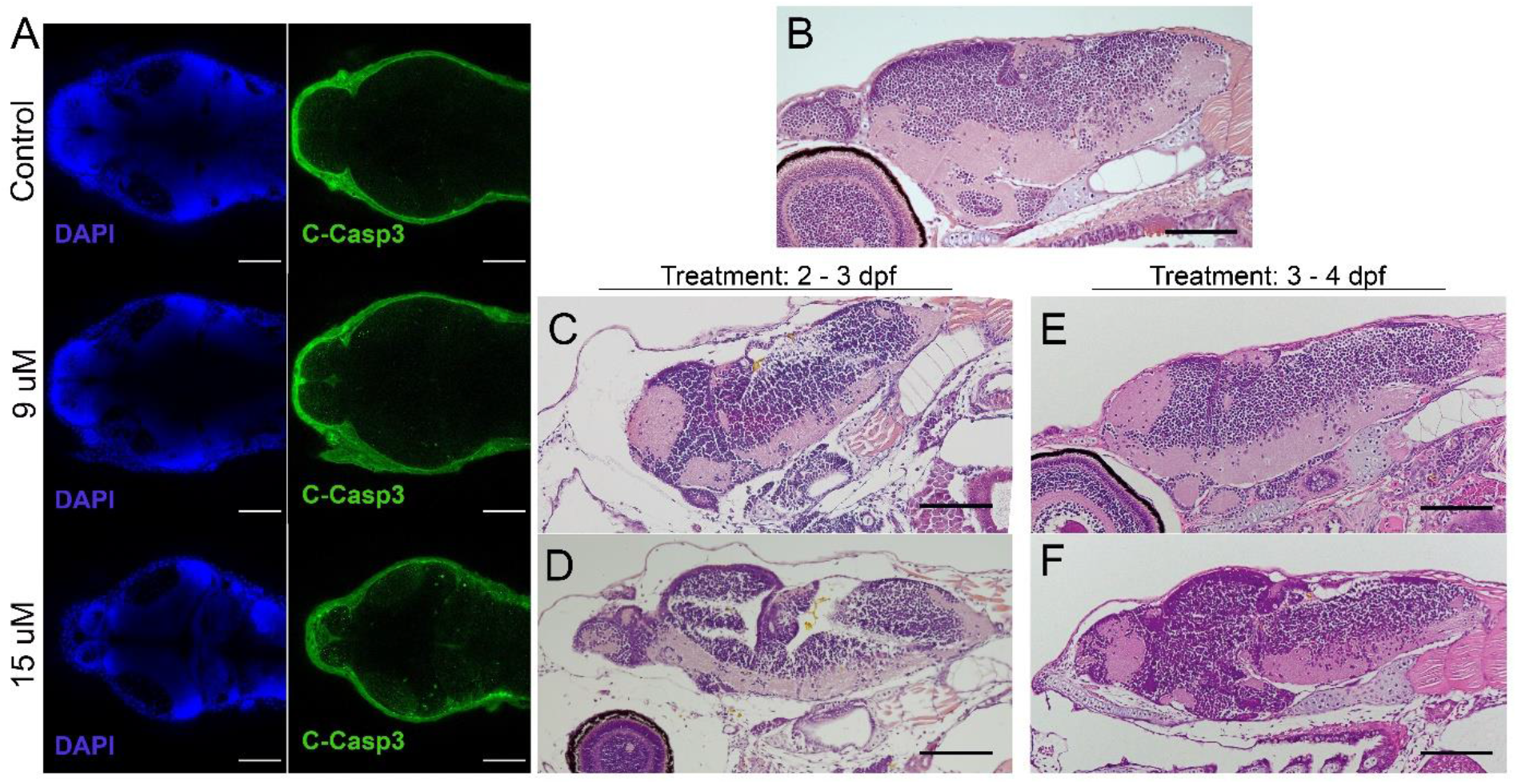
Bilirubin induced neurological damage (BIND) is detected in the larval brain: **A)** Apoptotic cells were detected in brains of control and hyperbilirubinemia larvae at 4 dpf. DAPI (blue) highlights brain structure, cleaved caspase 3 (C-Casp3) (green) labels apoptotic cells. **B-F)** H & E stained paraffin sections of 6 dpf larval brain. **A)** intact and healthy brain tissue in control larva; deformed brain tissue in larvae that received **B)** 9 μM, or **C)** 15 μM bilirubin between 2 - 3 dpf. **D)** Larva that received 9 μM bilirubin between 3 - 4 dpf had normal brain tissue. **E)** Larva that was exposed to 15 μM bilirubin between 3 - 4 dpf had slight deformation in brain tissue. Scale bars: 100 μm

### 3.5 Larval neurological functions are impaired in hyperbilirubinemia

In severe cases of hyperbilirubinemia, BIND may be presented as paralyzed or limited gaze, or weakness of eye muscles (24, 25). Eye reflexes of larvae with hyperbilirubinemia were measured at 6 dpf (2 days after washout). To this end, turn angle of eyes in response to light stimulus in semi-dark environment were measured and average turn angle was quantified (Fig. 7A, B). This experiment demonstrated that turn angle of the eyes decreased in a dose dependent manner.

**Fig. 7.**
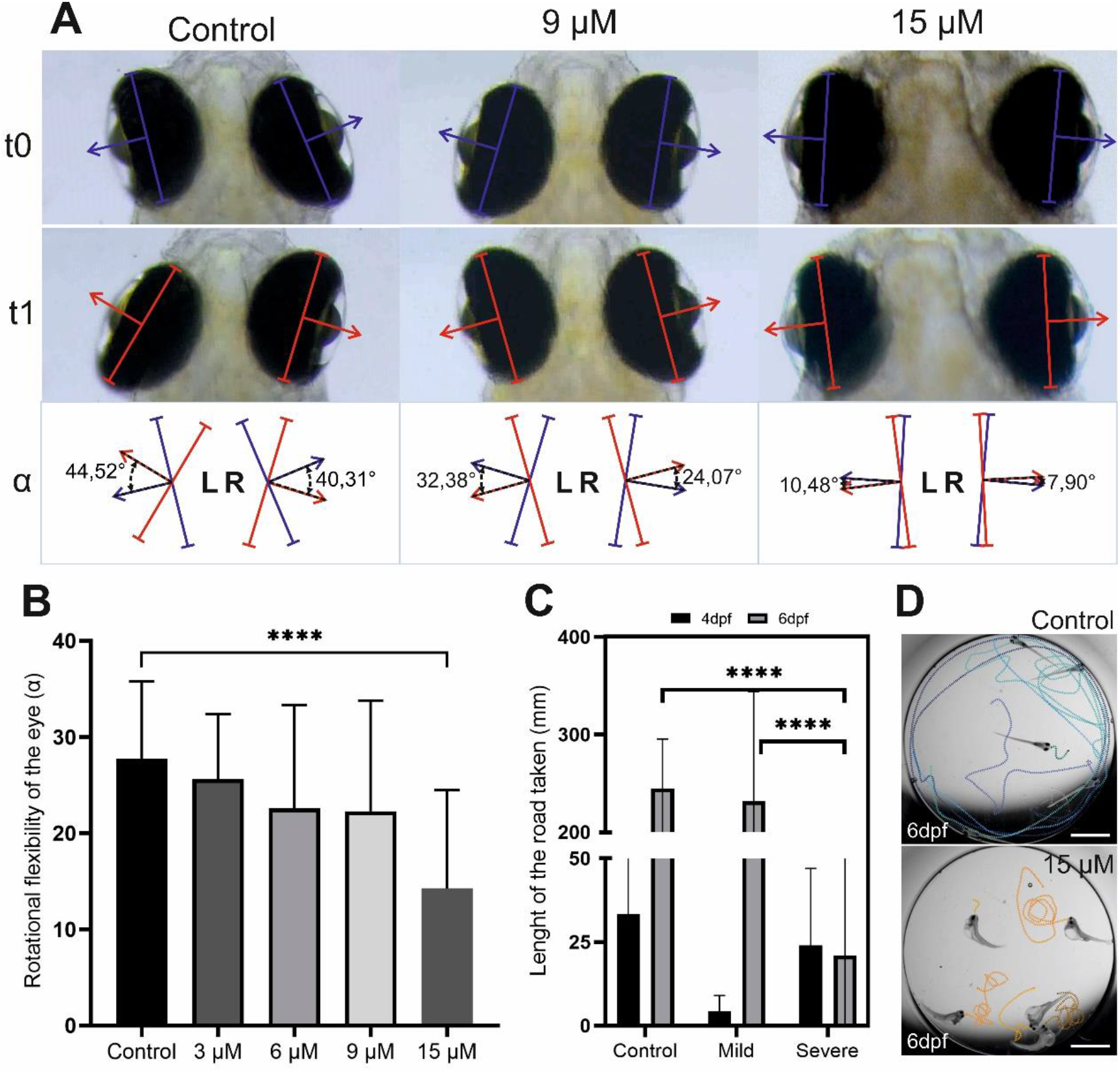
Hyperbilirubinemia affects eye reflexes and swim behavior: **A)** Turn of the eyes in response to light stimulant is displayed. Turn of the eye between t_0_ and t_1_ was measured as turn angle alpha. **B)** Average turn angle of the eyes of larvae exposed to 3, 6, 9 and 15 μM doses of bilirubin is displayed. **C)** Average swim distance of controls, and larvae exposed to 9 μM bilirubin between 2 – 3 dpf at 4 and 6 dpf is displayed. Data are Mean ± SEM. (A’’-C’’) n=20, 18, 22, 20 and 22 for control, 3, 6, 9 and 15 μM. **D)** Directional swim behavior of the control and swirling swim behavior of the severely affected larvae are displayed.

Motor dysfunction caused by BIND are dystonia (increased or decreased muscle tone) and excessive abnormal movements like writhing movements (26). The swimming behavior of the hyperbilirubinemic larvae with mild and severe phenotypes were recorded at 4 dpf and 6 dpf in order to test acute and lasting effects. Larvae with the mild phenotype initially displayed hypoactivity, swimming much less compared to controls at 4 dpf. However, at 6 dpf their activity levels increased and matched those of the control group (Fig 7C). Larvae with severe phenotype have curved body shape and they were not able to swim directionally but displayed a swirling type of swimming, at both 4 and 6 dpf (Fig. 7D). Compared to the control, these larvae were observed to display more activity and irritability (not quantified). At 6 dpf these severely affected larvae exhibited activity levels far below both the control group and larvae with a mild phenotype (Fig 7 C).

## 4. DISCUSSION

Although rodent models of hyperbilirubinemia and bilirubin induced neurological damage (BIND) have been developed before, alternative models are still needed to recapitulate newborn hyperbilirubinemia to the full extent (15). Here, the first zebrafish hyperbilirubinemia model is generated via exposure of early zebrafish larvae to external bilirubin. In order to mimic the human condition, the maturation of organs and expression of bilirubin metabolism enzymes during the first 5 days of zebrafish development were considered carefully. Since liver maturation is completed by day 5, and gut gains functionality by day 4, the period between 2 - 4 dpf was considered ideal for neonatal hyperbilirubinemia development (17, 27). One limitation of the study was insolubility of bilirubin in embryo water, which was overcome by use of a BSA carrier. After testing exposure between 2 - 3 dpf and 2 - 4 dpf, it was found that a 24-hour exposure between 2 - 3 dpf with 9 or 15 μM bilirubin induced a condition comparable to severe newborn hyperbilirubinemia. 15 μM caused a more severe phenotype and most larvae were highly damaged or dead by 5 dpf. Whereas 9 μM bilirubin caused a strong phenotype but better survival of the exposed larvae. Finally, 48-hour exposure caused stronger phenotypes in all concentrations, leading to lethality in 15 μM treated larvae, and severe phenotype in a small fraction of 6 μM exposed larvae. In sum, dose and time dependency of bilirubin accumulation and induced severe phenotype was observed.

Shifting the exposure onset time by one day, induced milder phenotype in overall larvae, suggesting tolerance to bilirubin increases already at 3 dpf. The tissues involved in bilirubin metabolism are liver, kidneys and gut in humans, but whether this is conserved in zebrafish was not studied previously (1). Here, tissue sections of bilirubin treated larvae of all groups were examined to check for bilirubin presence and any bilirubin induced damage. The fish that received bilirubin between 3 - 4 dpf had better morphology as well as healthier liver, and gut tissue at 6 dpf. The presence of bilirubin in these tissues 2 days after the washout suggests that the bilirubin that accumulated in the body are processed by liver and gut without damaging these tissues. On the other hand, fish that received bilirubin between 2 - 3 dpf had damaged liver and gut, although bilirubin was detected in these tissues as well. Damage to the liver and gut may be related to interference with the development of these tissues which is likely to hinder bilirubin clearance. This finding is important to show the conservation of bilirubin clearance mechanisms between human and zebrafish.

Total bilirubin retained in larval body was quantified with a spectrophotometric method and the amount of bilirubin in the body increased in a dose dependent manner in agreement with the phenotypic observations. Bilirubin accumulation in the brain was compared among groups that received different doses, and relative bilirubin accumulation in the brain reached a maximum in groups treated with 9 μM for 48 hours or 15 μM for 24 hours. The quantitation confirmed that 6μM bilirubin results in less accumulation in body and brain, compared to 9 and 15μM which are the doses that cannot be tolerated by 2 – 3 dpf zebrafish. This threshold of 9 μM can be considered similar to human significant hyperbilirubinemia with total serum bilirubin (TSB) ≥ 12 mg/dL (4). In human, total serum bilirubin level as well as the age of the infant are important indications for risk of BIND or kernicterus development, however these factors are not completely predictive. The reasons leading to individual variance in susceptibility to develop BIND or kernicterus is not fully understood. Zebrafish model exhibited a similar variance leading to development of mild or severe phenotype in equally treated and age matched zebrafish larvae can provide a means for studying the mechanisms underneath.

Next, BIND and related symptoms were investigated in the developed zebrafish model. Bilirubin accumulation in the brain and otic vesicle in a dose and time dependent manner was demonstrated. Bilirubin accumulated in telencephalon and rhombencephalon (cerebellum) in 9 μM exposed larvae, while increasing the dose to 15 μM resulted in intense accumulation in optic tectum (mesencephalon) as well. Similarly, in infants with bilirubin encephalopathy the hippocampus and ocular basal ganglia brain regions are affected (28). The brain compartments telencephalon, optic tectum, and cerebellum shrank and edema in the brain and eye sclera caused ballooning when toxic 9 - 15 μM doses were applied. The eye tissue was smaller indicating a delay in development, and the sclera width was increased. BIND in the zebrafish model was further confirmed by histopathology, and degeneration of the brain tissue was shown. Increase of apoptosis in the brain was also demonstrated with cleaved caspase 3 staining. When considered together, these findings indicate that zebrafish model can be used to induce BIND when 9 - 15 μM bilirubin is applied between 2 - 3 dpf.

Functional tests were performed to compare zebrafish phenotypes to human acute and chronic bilirubin encephalopathy (ABE and CBE). Curved body and deteriorated muscles as well as shorter body in zebrafish with severe phenotype, resembles opisthotonos that results from dramatic contraction of body muscles in severely affected infants. Problems with posture is generally accompanied with problems in movement control in affected infants. When swimming behavior was analyzed, it was found that these severely affected larvae responded to touch but were not able to swim directionally and displayed a swirling type of motion. These larvae had problems in terminating motion, but the swim distance was not increased due to the swirling swim style. On the other hand, larvae with mild phenotype had normal body posture and could perform directional swimming at 6 dpf, in a comparable manner to control group. Interestingly in the acute period at 4 dpf these larvae displayed decreased activity, lethargy. This phenotype is comparable to sleep tendency and increased lethargy observed in BIND patients (24). Upward gaze palsy is another symptom of bilirubin encephalopathy, reproduced in the zebrafish model as eye movement restriction phenotype (29). The turn angle of the eyes upon light stimulation decreased in 9 μM treated group and severely dropped in 15 μM treated group. It should be noted that the sclera widening also increases in 15 μM treated group which may be the main reason for gaze limitation.

## 5. CONCLUSIONS

Zebrafish newborn hyperbilirubinemia model reported here, showed that bilirubin exposure before development of the liver induces toxicity in a dose dependent manner. Larvae with hyperbilirubinemia were proved to be a good model for BIND. Not only the bilirubin accumulated in neural tissues but also it caused damage to the brain and the eyes, and induced lethargy. Strikingly, recovery of larvae from hyperbilirubinemia after washout was incomplete and even the mild cases (treated with 9 μM bilirubin) had defects in liver and brain tissues as well as some limitation of eye rotation at 6 dpf. Individual variability in susceptibility to bilirubin was observed, which can provide a basis for mechanistic studies. The model provides an alternative in vivo model for studying BIND mechanisms and prevention or treatment strategies.

